# Threshold illumination for non-invasive imaging of cells and tissues

**DOI:** 10.1101/2020.01.16.909911

**Authors:** M A Bashar Emon, Samantha Knoll, Umnia Doha, Danielle Baietto, Lauren Ladehoff, Mayandi Sivaguru, M Taher A Saif

## Abstract

Fluorescent microscopy employs monochromatic light which can affect the cells being observed. We reported earlier that fibroblasts relax their contractile force in response to green light of typical intensity. Here we show that such effects are independent of extracellular matrix and type of cell. In addition, we establish a threshold light that invokes minimal effect on cells. We cultured fibroblasts on soft 2D elastic hydrogels embedded with fluorescent beads to trace substrate deformation. The beads move towards cell center when cells contract, but they move away when cells relax. We use relaxation/contraction ratio, *λ*_r_, as a measure of cell response to light. The cells were exposed to green (wavelength, *λ* = 545-580 nm) and red (*λ* = 635-650 nm) light with a range of intensities. We find red light with intensity less than ~ 57 W/m^2^ results in *λ*_r_ = 1, i.e., cells maintain force homeostasis. Higher intensities and smaller wavelengths result in widespread force-relaxation in cells with *λ*_r_ > 1. We suggest the use of *λ* > 650 nm light with low intensity (*I* ≤ 57 W/m^2^) for time-lapse imaging of cells and tissues in order to avoid light-induced artifacts in experimental observations.

## 1. Introduction

Most living cells are photosensitive^1,2^. Furchgott and co-workers^3,4^ showed that smooth muscles of mammalian arteries under tonic contraction relax their force on exposure to light. Photorelaxation of fibroblasts was recently reported^5^. But light is essential for visualization. Since the advent of optical microscopy, various illumination techniques have been used to visualize cells and tissues. Recently however, monochromatic light is being used extensively for imaging. Many of these imaging methods, involving florescent reporters and green fluorescent protein (GFP), require blue and green light excitation with wavelengths less than 550 nm where photosensitivity cannot be ruled out, particularly if the cells are exposed to light multiple times for time-lapse imaging. A visible sign of photosensitivity or photo-toxicity is the change in cells’ morphology, such as blebbing, necrosis, formation of vacuoles, and mitochondrial swelling^2,6^. Such morphological changes may be expressed at different phases of cell division^7,8^. They may appear long after light exposure compared to the duration of exposure. We particularly observed change in traction force since it is essential for a variety of functions such as cell migration^9^, cellular homeostatsis^10^, differentiation^11,12^, morphogenesis, wound healing^13^, disease progression^14–16^. Our previous work showed that fibroblasts relax their forces partially when exposed to green light commonly used in florescent microscopes (wavelength, *λ* ~ 550 nm and intensity, *I* ~ 2250 W/m^2^) within 2 s of exposure^5,17^. We quantified the response by plating cells on soft Polyacrylamide (PA, 5 KPa) gel substrates functionalized with fibronectin and embedded with 100 nm fluorescent beads as fiduciary markers. The beads were placed within 1 μm from the surface of the gel substrate. Cells adhered to the substrate and generated contractile force. In response, the soft gel substrate deformed, and the beads followed the deformation, usually towards the center of the cell. The cell force reached a steady state within 3-4 hours of plating after which the net force became nearly stationary with negligible fluctuations over a short span of time (minutes)^18,19^. After reaching this steady state, we exposed the cells with regular green light for either 2 s or 60 s, while we monitored the dynamics of the beads^5,17^. We found that soon after light exposure: (1) the beads moved mostly away from the cell-center implying force relaxation, while a minority of beads moved towards cell-center implying contractility. This suggests a net force reduction, i.e., photorelaxation; and (2) some of the beads moved abruptly outward (“jumping”), implying sudden local force relaxation^20^.

Here, we seek to establish a threshold light intensity and wavelength that do not affect the cells, i.e. the cells maintain force homeostasis without photo-relaxation, and yet the light is sufficient for high-resolution time-lapse fluorescence imaging. We performed our experiments by exposing fibroblast (monkey kidney, CV-1 cell line) cells to green (*λ* = 545-580 nm) and red light (*λ* = 635-650 nm) with a range of intensities. The cells were plated on PA gel substrate functionalized with fibronectin and embedded with florescent beads. We measured the motion of beads for 1 hour using a timelapse imaging approach. We quantify the ratio *λ_r_* of net bead displacement away from and towards the cell center. *λ_r_* > 1 implies that the cell has relaxed, *λ_r_* = 1 implies that the cell is maintaining force homeostasis. We found *λ_r_* = 1 for red lights with intensity of *I* = 57 W/m^2^ or lower. For higher intensities and lower wavelengths, we find *λ_r_* > 1. We also quantified the net contractile force of the cells, which also remains unaffected by the red light. Above the threshold intensity, cell contractile force decreases with time. This threshold is not only limited to cells plated on fibronectin as Extra-cellular matrix (ECM). We verified that the threshold also applies to cells cultured on laminin-coated substrates.

## 2. Materials and Methods

### 2.1 Cell culture and substrate preparation

We used three types of cell lines for the study: normal monkey kidney fibroblast, CV-1 (ATCC), normal human colon fibroblast, CCD112CoN (ATCC), and mouse embryonic fibroblast, NIH 3T3 (ATCC). The effect of light was studied for both 2D and 3D culture. CV-1 and CCD112CoN were cultured on 2D substrates, 3T3 were cultured in 3D. Inspected cells on 2D substrates were more than 100 μm away from neighboring cells and the cell density in 3D collagen culture was approximately 0.1 million/ml. CV-1 cells were plated on Polyacrylamide (PA) hydrogels embedded with fluorescent particles. The beads were localized at a depth of about 1 μm from the top surface of the substrate following a protocol outlined by Knoll^21^. We used two types of fluorescent beads- i) 0.2 μm-dia dark red beads (excitation/emission-660/680 nm, Thermo-Fisher) ii) 0.1 μm red beads (excitation/emission-580/605 nm, Thermo-Fisher). The elastic modulus of the hydrogels was approx. 5 kPa (4.47 ±1.19) based on the protocol established by Tse and Engler^22^. Substrates were functionalized by two types of ECM, namely fibronectin (Human, Corning) and laminin (Human, Corning). Two ECMs were chosen to eliminate the possibility that photo-relaxation might be an artifact of a particular ECM. The protocol for functionalization is outlined by Tse and Engler^18^. Briefly, 0.2 mg/ml sufosuccinimidyl-6-(4’-azido-2’-nitrophenylamino)-hexanoate (Sulfo-SANPAH, Thermo Scientific) solution in HEPES buffer (50 mM HEPES at pH 8.5, Fisher Scientific) was applied to the PA gels and then was activated with UV. The substrates were then immersed overnight in fibronectin or laminin solution in HEPES buffer. Concentration of 25 μg/ml was used for both fibronectin and laminin. The gels were then washed with PBS and were ready for cell culture.

CCD112CoN (ATCC) cells were tested on 2D PA gel substrate following the above protocol. However, only fibronectin functionalization was tested.

NIH/3T3 (ATCC) cells were mixed with rat tail collagen type I (Corning) such that final concentration of collagen was 2.5 mg/ml, and cell density was 0.1 M/ml in cell-collagen mixture. This mixture was dispensed in a glass-bottom petri-dish (well diameter 12mm, depth 1 mm, Cellvis). After filling the well with cell-collagen mixture, the well was covered with coverslip (22mm, EMS). The petri-dish was then flipped and immediately put in the refrigerator at 4 °C so that the cells settle down to the cover slip while the collagen does not completely polymerize. After 15 minutes, samples were flipped again and kept in the incubator (37 °C) for 3 minutes which allowed the cells to fall under gravity through the liquid collagen while it polymerizes simultaneously. The petri-dish was flipped 2 more times and the time intervals between the flips were 5 and 7 minutes respectively in the incubator. The cover slip was removed and the dish was filled with cell-culture media. From z-stack images, sample thickness was found to be 900 microns and cells were observed in different planes. We selected a plane 500 μm above the bottom of the dish to ensure that cells were completely surrounded by 3D collagen matrix. Cells for all of above experiments were cultured in fibroblast media containing 90% Dulbecco’s modified Eagle’s medium (DMEM, Corning), 10% fetal bovine serum (FBS, Gibco and 1% penicillin–streptomycin (Lonza).

### 2.2 Light source and illumination methods

The cells were illuminated continuously with either: (1) a deep red collimated LED (light emitting diode) (Thorlabs, Inc., Newton, NJ) coupled with far red filter set (Semrock Brightline LF635/LP-B-000, Rochester, NY) giving λ = 635-650 nm considering full width at half maximum (HMFW) intensity or (2) a fluorescent metal halide lamp (X-Cite®Series 120, Excelitas Technologies, Waltham, MA) coupled with an mCherry filter (Semrock Brightline mCherry-M-OMF, Rochester, NY) giving λ = 545-580 (HMFW). These range of wavelengths were measured by spectral analysis of the illumination using a standard spectrometer (USB2000+, OceanOptics) (Suppl. Fig. 1). Technical details about the light sources and the filters are provided in Supplementary Information. Suppl. Fig. 2 shows a schematic diagram of the experimental setup and light paths. Neutral density (ND) filters were employed for both sources to tune light intensity. For example, ND100 and ND25 indicate 100% and 25% of the light were transmitted. In addition to ND filter, power output from the fluorescent metal halide lamp for mCherry light was modulated to control light intensity. Thus, 25%ND25 means that 25% of light was allowed from the source, and 25% of this output was filtered by the ND25 filter. The intensity of LED light was controlled by the ND filters only. The light sources and their intensities are shown in Table 1. Light intensities were measured using 40X water immersion objective (Numerical Aperture, NA=1.15, Olympus) by PM100 power meter (ThorLabs) at a plane 15.5 mm above the objective and at the focal plane which is the typical location of the sample. Note that the power meter aperture is large (9.5 mm) and could not be used to quantify the special intensity profile of the incident light. In order to reduce the aperture, the sensor was wrapped with an aluminum foil with 0.75 mm diameter aperture. Thus, the intensity was measured for light passing through the small aperture only. The power meter was moved orthogonal to the incident light and along the radial direction of the light beam to measure the spatial intensity profile. Also, it was not possible to measure light intensity at the sample planes during experiments due to the presence of cells. Hence intensity was measured at a distance of 15.5 mm above the objective. Although, the intensity can then be estimated at the focal plane (sample plane) from the measurements at the 15.5 mm plane, we measured the intensity at the focal plane as well *in the absence of cells* to ensure mimimal loss of light due to dispersion. The correlation between the measured intensities at the two planes was used to estimate light intensity on cells. Intensity profiles of representative light sources are provided in Suppl. Fig. 3.

**Table 1.** Particle tracking precision for various illumination sources. Displacement distributions for cells exposed to various light sources, listed in order of increasing light intensity. Particle tracking precision determined by tracking beads immobilized in PA gels devoid of cells during one-minute of continuous illumination through 40X water immersion objective (NA=1.15). Illumination source column indicates light source (LED or mCherry), as well as the accompanying neutral density (ND) filter classification. ND100 indicates no filter was used (i.e. 100% of light passed through). Modulation of relative power output for the fluorescent metal halide lamp utilized as the mCherry source is indicated, preceding the ND classification (i.e. 12 or 25%). Tracking precision computed as standard deviation of the Gauss fit of the displacement distribution for each illumination source. Each illumination source represents over 1,000 particles from 3 distinct gel substrates. * Values reported was determined from extrapolation of experimental data. ** Sample plane intensity was determined employing intensity profiles in suppl. Fig. 3.

| Lighting Condition | Avg. intensity at 15.5 mm from the objective ($\text{W/m}^2$) | Max. intensity at cell sample plane ($\text{W/m}^2$)** | Measurement Resolution (nm) |
| --- | --- | --- | --- |
| LED ND6 | 0.12 | 14.6 | 18.07 |
| LED ND12 | 0.28 | 34.1 | 14.84 |
| LED ND25 | 0.47 | 57.2 | 10.62 |
| LED ND100 | 1.93 | 234.8 | 6.1 |
| mCherry 12% ND6 | 1.43 | 257.7 | 13.11 |
| mCherry 25% ND6 | 2.47 | 445.0 | 9.77 |
| mCherry 12% ND12 | 3.17 | 571.1 | 9.05 |
| mCherry 25% ND12 | 5.53 | 996.4 | 7.57 |
| mCherry 25% ND25 | 12.5 | 2252.2 | 11.24 |
| mCherry 25% ND100 | 40.8 | 7351.1 | 6.71 |
| mCherry 100% ND25 | 41.65* | 7504.2 | - |
| mCherry 100% ND100 | 135.94* | 24493.7 | - |

### 2.3 Particle tracking and limitations

The motion of fluorescent beads was tracked from images for analyzing their movements with time. First, the cell spreading area was determined and the centroid of the area was established using the following formula: centroid 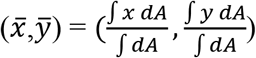. Let the vector from the origin (cell centroid) to an embedded florescent bead be *<u>r</u>*(*0*) and *<u>r</u>*(*t*) at *time* = *0* and *time* = *t* respectively. Hence, the net magnitude and direction of displacement of the bead is given by |*<u>r</u>*(*t*)| – |*<u>r</u>*(*0*)|. Motions of all the beads within cell area were computed and statistically analysed for cells subjected to the described illumination protocols. Motion toward the centroid (inward) is negative and represents cell contraction; motion away from the centroid (outward) is positive and represents cell relaxation. Relaxation was assessed by a majority of motion outward relative to inward. It is expected that cells exposed to sufficiently low intensity light would not exhibit light-induced relaxation. However, lowering the excitation light intensity reduces the signal level with respect to background noise, reducing the particle tracking precision. We quantified the noise and measurement resolution for each light source (see Table 1) by tracking fluorescent beads in PA gels without adherent cells. For each illumination condition, light was shined for 60 s. Bead displacement was quantified as the change of position with an arbitrary origin during 60 s, i.e., |*<u>r</u>*(*60*)| – |*<u>r</u>*(*0*)|. A Gaussian curve was fit to probability distributions of the particle motion. The standard deviation (*σ*) of the distribution represents a measure of displacement resolution for each light source (Table 1, Figure 1). For evaluation of cell response, sparsely populated fibroblasts were then cultured on the same substrates with beads. Single cells were then exposed to various illuminations (Figure 1) for prescribed durations. The displacements of beads within the footprint of each cell were then analysed and reported.

**Figure 1.**
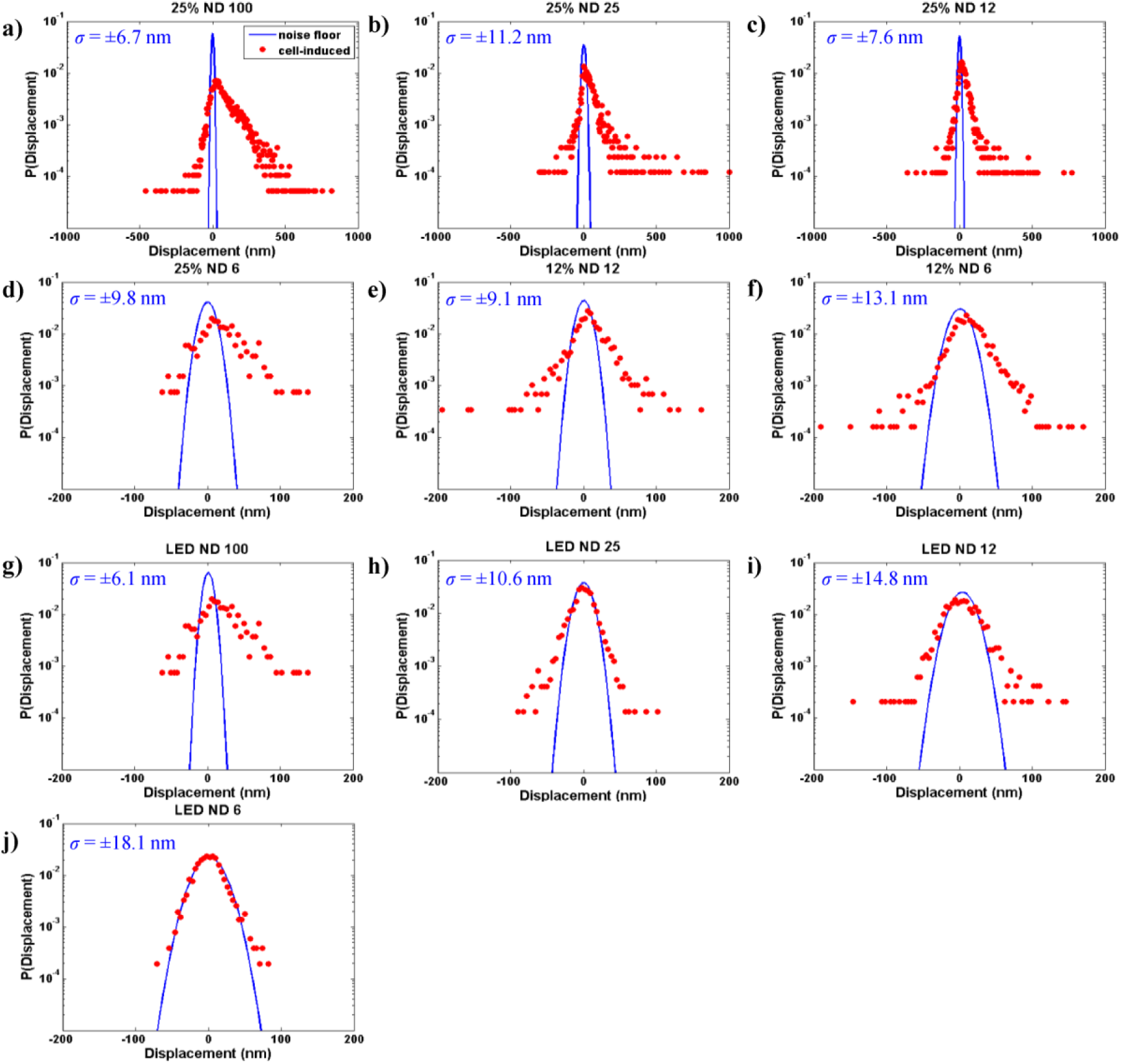
Determination of noise and signal. Probability distributions of particle (florescent beads) displacements embedded in PA gel substrate with (red) and without (blue) cells at various illumination conditions (a-j). For each case, light was shined for 60 s. Particle displacement is quantified as the change of position during 60 s, i.e., **<u>r</u>(60)** – **<u>r(0)</u>**|, where **<u>r</u>(t)** is the vector location of the particle. Tracking precision is estimated as the standard deviation of Gauss fit of the displacement distribution for the cell free substrates. Each illumination source represents over 1,000 beads from 3 distinct gel substrates. The variances of the distributions were determined according to an F-test (α = 0.05).

## 3. Results and Discussion

### 3.1 Effect of light on cells to monitor florescent beads embedded in substrates

One of our objectives was to monitor cell traction over long times after the cells are exposed to light. But monitoring also requires light to fluoresce the tracer beads embedded in the substrate. In order to minimize the effect of monitoring light on cells, we use dark red beads that can be fluoresced with sub-threshold light (long wavelength LED, ex/em-660/680 nm with I < 57 W/m^2^). However, when monitoring is done while the cells are exposed to 60 or 120 s mCherry light, we use 0.1 μm red beads (ex/em-580/605 nm). Thus, no additional exposure is needed for monitoring. Similarly, when beads are monitored while cells are exposed to continuous LED light, we used dark red beads (ex/em-660/680 nm). When beads are monitored after cells are exposed to prescribed target dosage of light, the monitoring illumination with sub-threshold light may still affect cell traction. To address this issue, we carried out an extensive dosage analysis for various illumination conditions (Table S1). The analysis reveals that for most of the cell exposure conditions, initial target dosage on cells for 60 s or 120 s is more than 96% of the total illumination which indicates that the effect of light for monitoring is minimal.

### 3.2 Effect of various light dosages on cells

In order to evaluate the effect of illumination on cells, we consider two options: (1) reducing the amount of radiation energy to the cells, and (2) reducing the amount of time the cells are exposed to a given light intensity.

For Option (1), we note that adherent, non-motile cells maintain a steady contractile state, or force homeostasis after a few hours of plating^6,18,23–26^. Thus, we search for the light (wavelength and intensity) that allows cells to maintain homeostasis. We exposed CV-1 cells, plated on PA gels functionalized with fibronectin and laminin independently, to continuous illumination for 60 s with various lighting sources. Substrate deformations were assessed as “contractile” when the beads moved towards the cell center, or “relaxation” when the beads moved outward^21^. We measured cell contraction or relaxation using the LED light source (λ = 635-650 nm with I < 57 W/m^2^) for the following 1 hr. For Option (2), we exposed the cells to green and red lights for 15 s and 2 s and measured relaxation using LED for the following 1 hr. Our findings are presented in the following sections.

### 3.3 Lower wavelength light causes more photo-relaxation

Probability distributions (normalized histograms such that the area under the histogram equals to 1) of displacements induced by cells exposed to each light source are shown in Fig. 1a-j (red points). Gauss-fit of the noise floor is superimposed on the histograms (blue lines in Fig. 1a-j, merged plot shown in Suppl. Fig. 4). Deviation of the cell-induced motion (red) from the noise-floor (blue) indicates the extent to which active cell motion occurs beyond the measurement noise. The variances of the distributions were determined according to an F-test (α= 0.05). A positive skewness indicates relaxation whereas a negative skewness means contraction. Substrate deformations of all mCherry-exposed cells exhibit significant statistical deviation from the noise-floor (Fig. 1a-f). This also holds true for cells exposed to the LED ND100 source (Fig. 1g). However, for LED ND6, a large portion of the cell-induced motion aligns with the noise floor (Fig. 1j). Thus, very little activity is detected beyond the measurement noise for LED ND6 source. This can be due to the fact that the cells do not produce any excess force on the substrate during this subsequent 60 s period. For LED ND25 and ND12, cell induced motions are detected beyond noise floor, and the distributions are symmetric, i.e., the cells’ contractility and relaxation during 60 s time are balanced, resulting in force homeostasis. While cells can maintain force homeostasis under exposure to any of the sources (LED ND25, LED ND12, LED ND6), we identify LED ND25 as a threshold light source that offers a compromise between minimal photo-relaxation and sufficient brightness for time-lapse imaging of cells and substrate beads.

We evaluate the probabilities of contraction, (*P_c_*) and relaxation (*P_r_*) by integrating the area under the left and the right sides of probability distributions (red points) in Fig 1 for each illumination condition. We then evaluate the relaxation ratio, *λ_r_* = *P_r_*/*P_c_* The probabilities of relaxation compared to contraction increases with light intensity and lowering wavelength (Fig. 2a). This suggests that wavelength may also play an intrinsic role, as higher wavelength sources are known to reduce damage to cells^27^. Cells exposed to light with intensity below 57 W/m^2^ exhibit equal proportions of contraction and relaxation with *λ_r_* ~ *1.0* (Fig 2a). We found a correlation (Fig. 2b) between relaxation/contraction ratio (*λ_r_*) and illumination intensity by best fitting *λ_r_* to a set of experimental data (green) that involve all of the light intensities. We then used a separate set of data (orange) to validate the correlation. *λ_r_* for CV-1 cells with light intensity *I* (W/m^2^) can be estimated by the following equation:

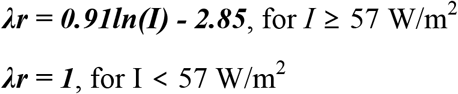

**Figure 2.**
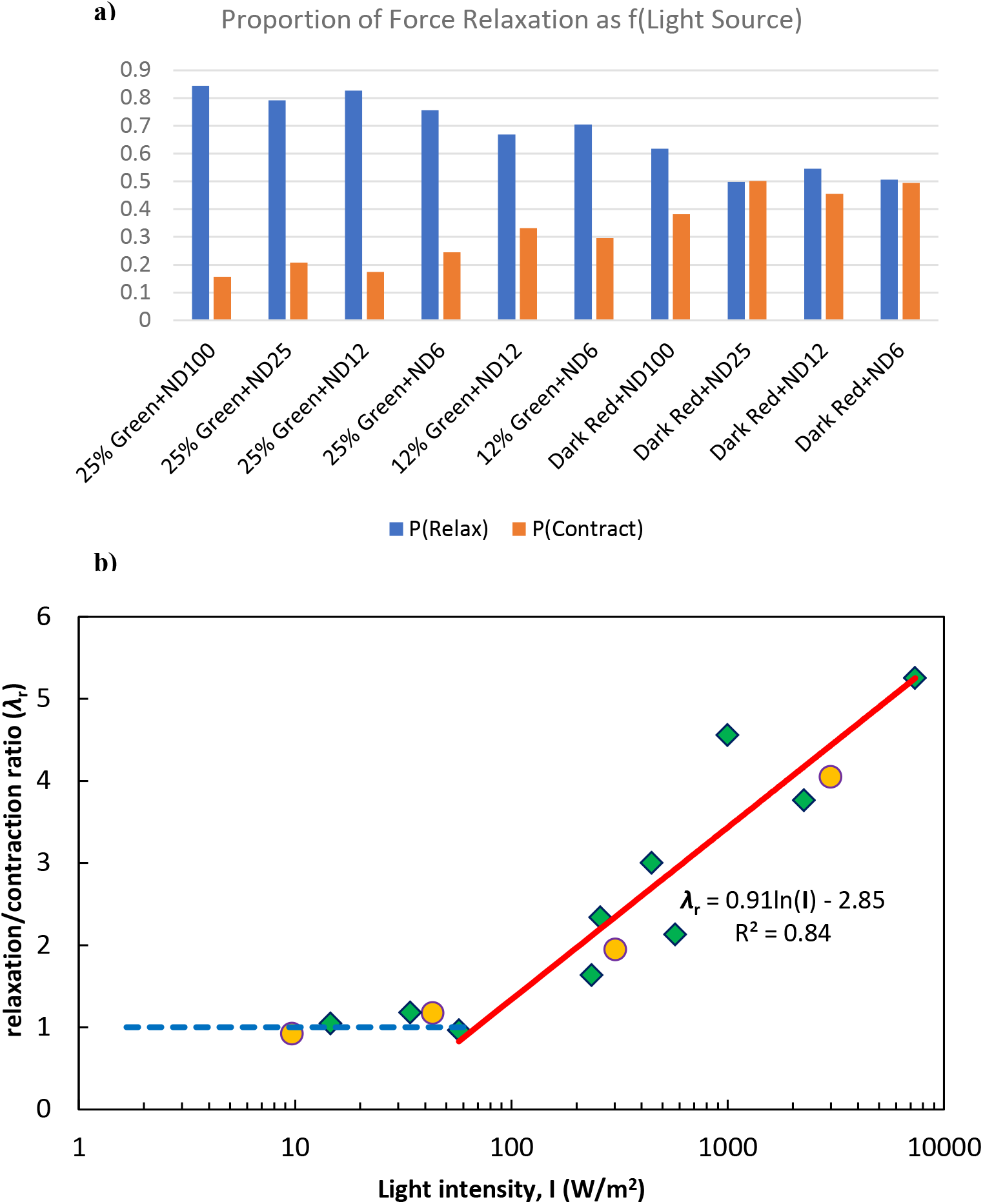
a) Cell force relaxation decreases with decreasing light intensity during illumination. Proportion of outward-relative to inward-moving beads represents decreasing dominance of force relaxation over contraction throughout illumination. Probability of outward-(P(Relax)) moving bead displacements relative to the cell centroid decreases with decreasing illumination intensity, where the probability of inward motion, P(Contract)=1-P(Relax). All cells were illuminated continuously for 60 s. Each illumination condition represents n=5 distinct cells. **b) Relaxation/contraction ratio (λ_r_) has a logarithmic relationship with illumination intensity.** Above I=57 W/m^2^, λ_r_ can be predicted by the red line with equation λ_r_= 0.91ln(I) - 2.85. Below I=57 W/m^2^, we expect to have no relaxation and hence λ_r_ = 1. Curve in the inset shows the same relation on a normal scale. Data points with green markers were used to generate the correlation. Orange markers are the data points used for verification of the relation.

### 3.4 Time evolution of traction forces suggests that cells maintain a steady state following initial illumination period

Traction forces of living cells have traditionally been thought to attain a steady state soon after (~ 2 hrs) adhering to a surface^6,18,23–26^. Cells build up traction force over minutes to hours of adhesion and maintain that force until initiating other activities (i.e. migration or division). In spite of our finding that light exposure affects cell forces, we aimed to understand how traction forces evolve over time following illumination with potentially damaging lighting conditions. We expect that traction forces would decrease continuously over time following exposure to relaxation-inducing light, until the cell completely detaches from its substrate. We also expect that the rate of traction force reduction would be proportional to the illumination dose (intensity and illumination time), as indicated by prior results highlighting a dose-dependent response^21^.

For option 2 (reducing exposure time), four different test lighting sources were utilized: mCherry 25% ND100 for (1) t=15 s and (2) t=2 s, and LED ND100 for (3) t=15 s and (4) t=2 s. The LED ND25 was chosen as the non-damaging light source for traction monitoring (Table 1, Fig. 1). As expected, the greatest decrease in traction force occurs for the mCherry 25% ND100 for t=15 s condition, followed by mCherry 25% ND100 for t=2 s, LED ND100 for t=15 s, and LED ND100 for t=2 s (Fig. 3). The traction forces for all sources decrease initially within first 15 min, with mCherry 25% ND100 t=15 s and t=2 s decreasing the most, followed by LED ND100 t=15 s and t=2 s, respectively (Fig. 3). Following the initial decrease, the force values for each light condition maintain a steady-state, fluctuating only ±5%, for the next one hour. However, force relaxation decreases with decreasing light intensity. In contrast to net force relaxation, the cells appear to maintain a steady ratio of contraction and relaxation after short light exposure. A measure of the change in contraction-relaxation is given by the probabilities of bead displacements inward (contraction), *P_c_*, and outward (relaxation), *P_r_*, of the cell center (demonstrated in Fig. 2a) respectively. These probabilities are shown in Fig. 4 for cells exposed to mCherry 25% ND100 for t=2 s, LED ND100 for t=15 s, and t=2 s. Soon after exposure to mCherry 25% ND100 for t=2 s, *λ_r_* = *P_r_*/*P_c_* ~ *0.7*. This ratio increases slightly over 60 mins even though net force decreased by 20% during this time (Fig. 4). *λ_r_* is close to 0.5 soon after exposure to LED ND100 light and increases slightly during the following 60 mins.

**Figure 3.**
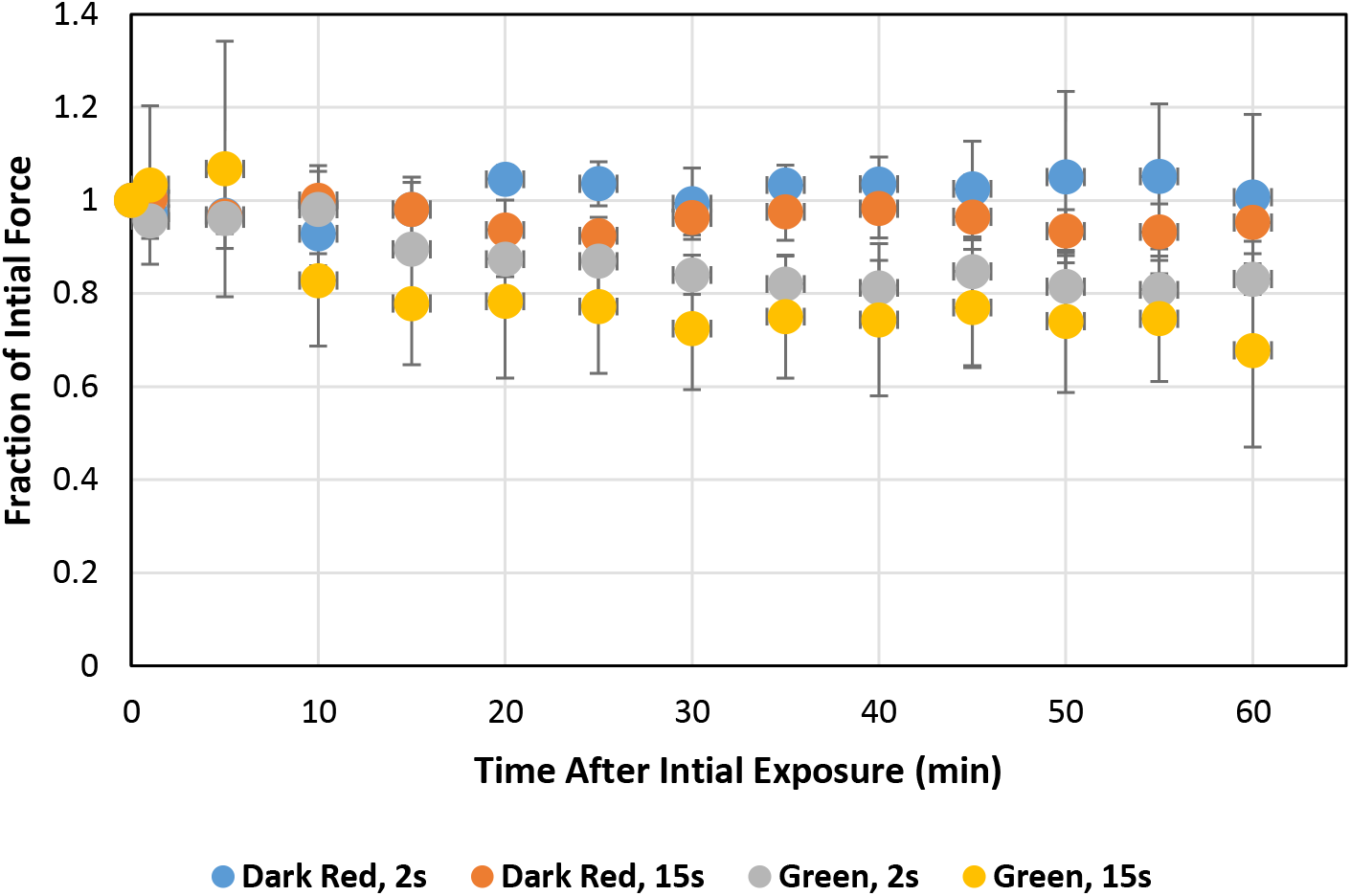
Time evolution of traction forces as a percentage of initial force. Traction force over time for cells exposed to any of the following illumination sources: LED (Dark red light) ND100 for t=2 s (blue), LED (Dark red light) ND100 for t=15 s (orange), mCherry 25% (Green light) ND25 for t=2 s (gray), or mCherry 25% (Green light) ND25 for t=15 s (yellow). Traction force listed as a function of initial force prior to illumination. Time t=0 represents the force at the instant the illumination period ended. Each distribution represents an average for n=3 cells. Error bars represent standard deviation.

**Figure 4.**
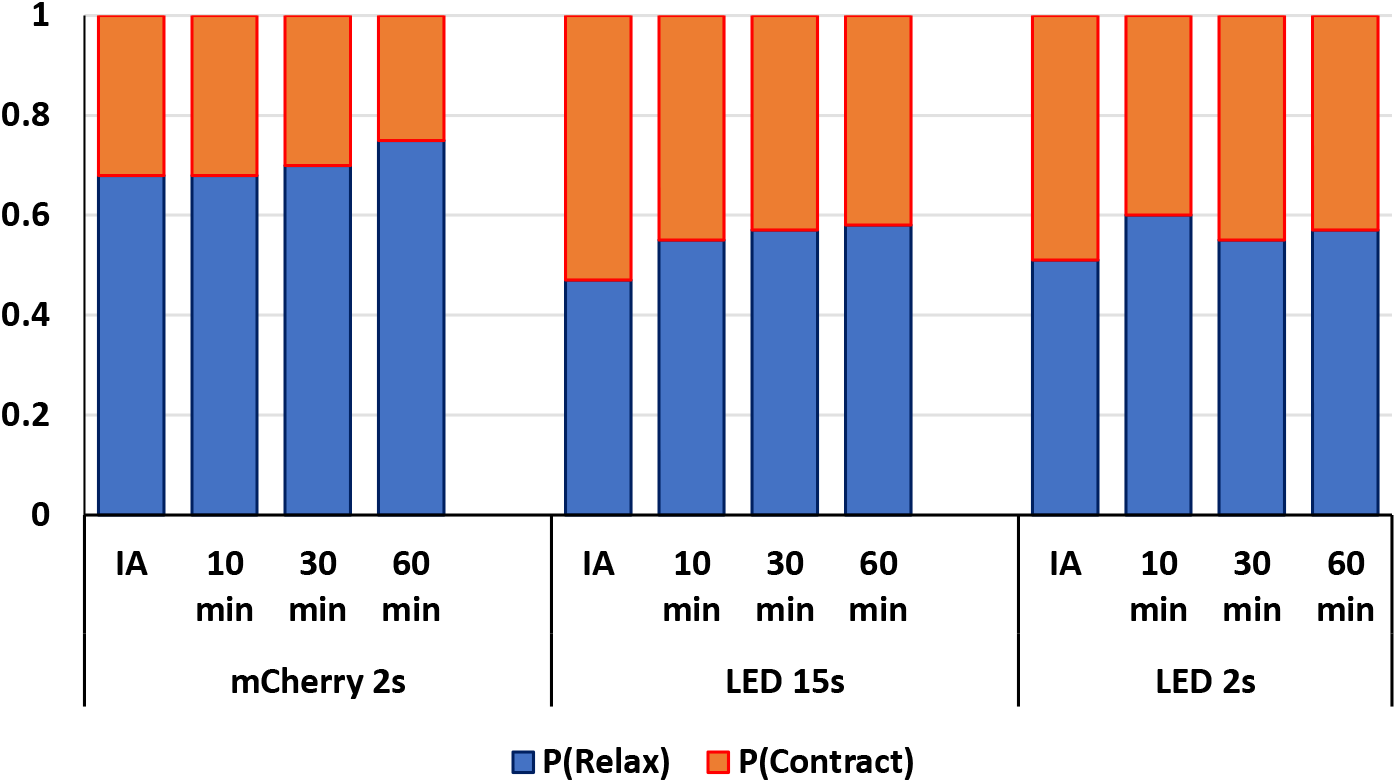
Proportion of force relaxation maintained over time following illumination for various exposure conditions. Probability of relaxation (P(Relax)) and contraction (P(Contract)) determined as the proportion of outward-versus inward-moving particles for each illumination condition (mCherry 25% ND100, t=2 s; LED ND100, t=15 s; LED ND100, t=2 s). Force relaxation shown at 0 s (Immediately after, IA), 10 min, 30 min, 60 min after illumination, all normalized with the force at 0 s. Each curve represents an average of n=3 cells and number of beads >1000.

### 3.5 Evidence of localized force changes over time

Even though cells maintain *λ_r_* over time, inspection of the individual bead displacements arising from local traction forces yields evidence of small, localized force changes. Heat maps of cell-induced displacements reveal modulation of force application strength throughout the spatial area of the cell (Fig. 5). While the location of the largest displacements shifts throughout the one-hour observation period, the dominant angle/orientation along which the forces are aligned appears to remain constant.

**Figure 5.**
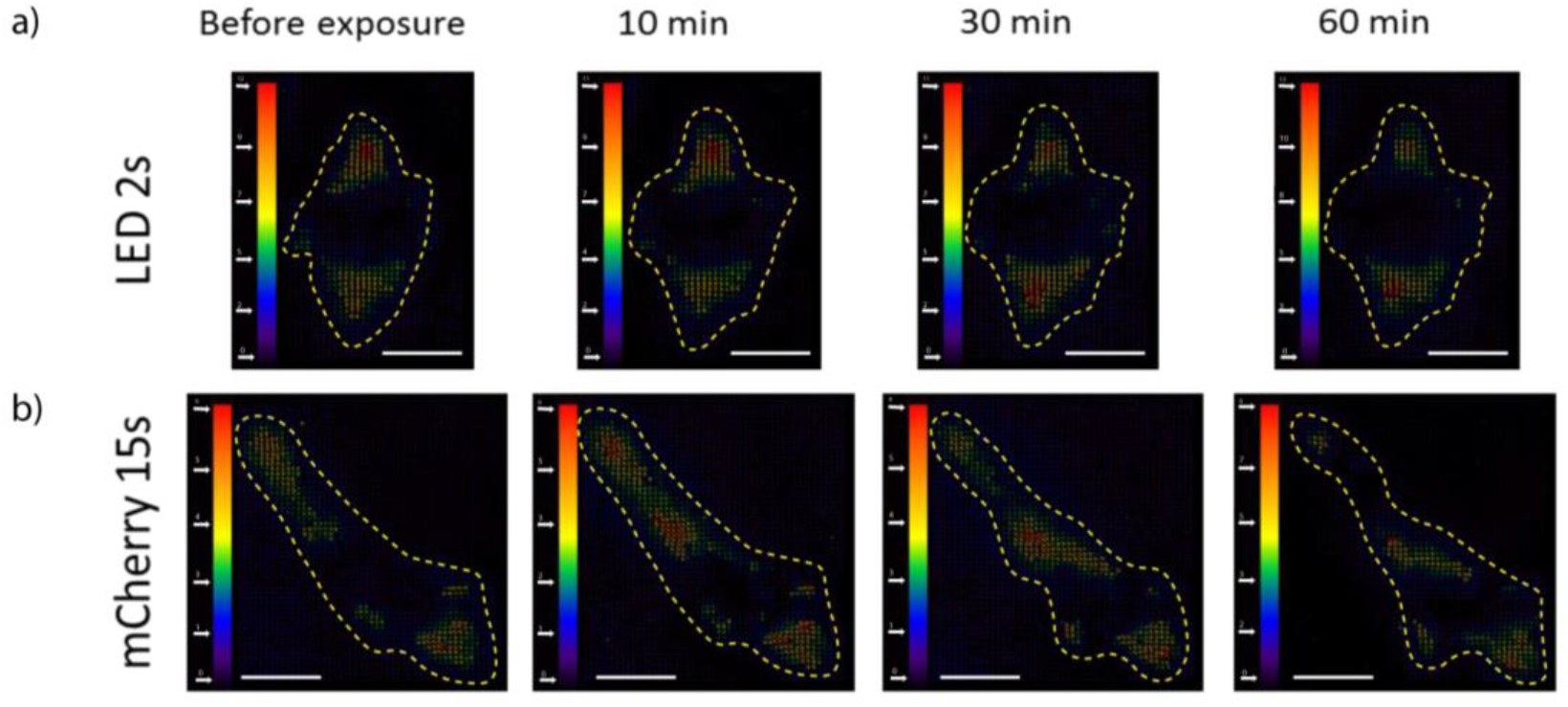
Displacement maps exhibit steady state traction force following illumination. Maps of displacement as a result of cell traction following illumination by either **a)** LED ND100 for t=2 s or **b)** mCherry 25% ND100 for t=15 s. Unit bar in pixels. Scale bar: 20 um.

### 3.6 Photo-relaxation is independent of ECM functionalization

In order to test whether the observed photo-relaxation is an artifact of fibronectin functionalization, we plated CV-1 cells on gel substrates (*E* = 5 KPa) functionalized with laminin. We exposed the cells to the following lighting conditions: a) LED ND25 (I= 57W/m^2^), 120s b) mCherry 100% (green light) with ND25 (I=7500 W/m^2^) and ND100 (I=24500 W/m^2^ seems too high?), each for 120 s. For all light conditions, the beads were imaged every 5 s for a total of 2 mins. There was no significant bead movement/force relaxation within this period for LED ND25, 120s light which agrees with the experimental results on fibronectin-coated substrates. Also, this ensured suitability of LED ND25 illumination for time-lapse imaging to investigate the effect of mCherry light. Probability distribution of bead displacements from the experiment with mCherry is presented in Fig. 6. Both distributions are positively skewed which indicates force relaxation due to both illuminations. Also, as expected, the distribution was relatively more skewed for 100% mCherry ND100 due to higher intensity. Corresponding bead displacements for representative cells over a 120 s period is shown in Suppl. Fig. 5 A-B. It is clear that force relaxation is significantly more pronounced as compared to contraction, which is similar to the results with fibronectin. These results suggest that photo-relaxation is independent of ECM, and LED ND25 with intensity 57 W/m^2^ is a safe threshold for time-lapse imaging for both laminin and fibronectin coated substrates.

**Figure 6.**
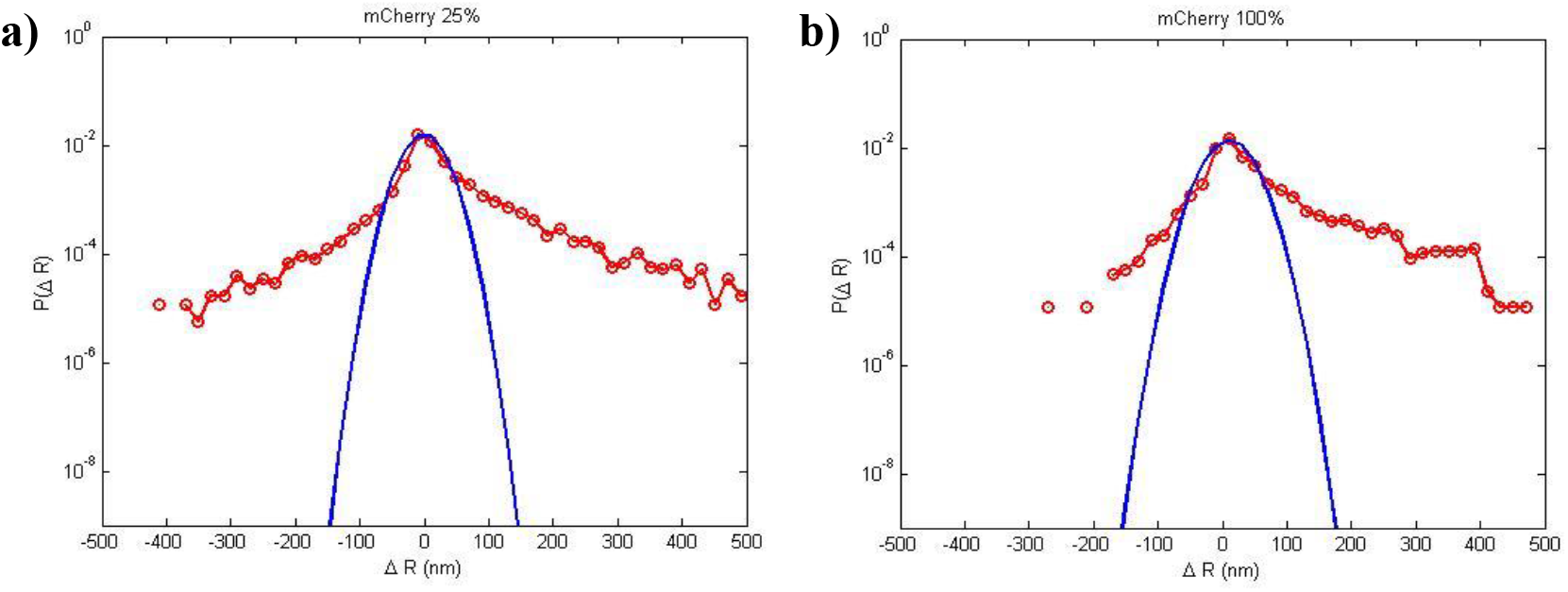
Light exposure induces force relaxation in cells on Laminin-coated PA gel substrates. Probability Distribution Functions of bead displacements within cell footprint for CV-1 cells. The substrate is functionalized with Laminin ECM. The cells are exposed to 100% mCherry (green) light with either **a)** ND25 or **b)** ND100 Neutral Density filter. ND100 implies no filter. Both the plots are positively skewed, indicating more positive (outward) bead motion, caused by ‘force relaxation’ of the cells during 60s of continuous illumination.

### 3.7 Photo-relaxation affects multiple cell types across different species

We anticipated that photo-relaxation is not limited to CV-1. To this end, we explored the effect of light on a human (CCD-112 CoN, colon normal fibroblast) and mouse cells (NIH/3T3 fibroblasts); although majority of our study was carried out with CV-1 cell line. We performed traction force microscopy on CCD112CoN cells during and after 120s exposure to i) LED ND25 presented in Fig. 7. As expected, both mCherry lights induced force relaxation throughout the entire 120 s of illumination. During the following hour, these cells maintained a steady state as observed with CV-1 cells as well. On the other hand, the LED light source, which illuminated at an intensity level below the established threshold, triggered no such relaxation. In fact, at the end of the illumination period, mean force seems to be higher than the initial value, although over the subsequent hour it fluctuated and reduced by a small amount. Presumably this reduction is due to natural variation since no instantaneous response was noted at the exposure which is common in cases of high intensity. Hence, we can deduce that human fibroblasts are also susceptible to photo-relaxation if irradiated with above-threshold light. However, the precise threshold light for CCD112CoN may differ from that for CV-1 cells.

**Figure 7.**
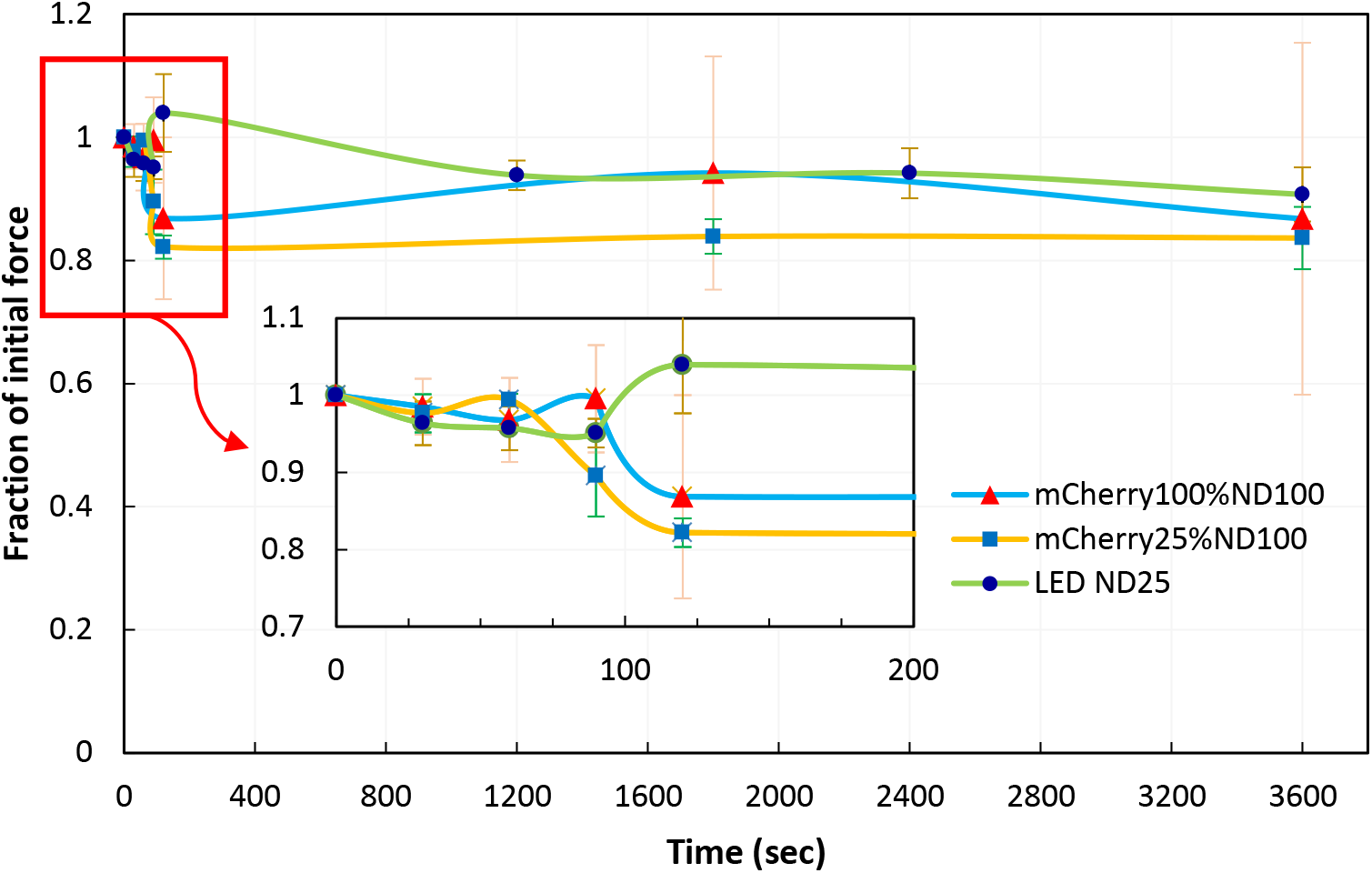
Evolution of traction as a percentage of initial value for CCD112CoN cells. Traction force measured over time for cells exposed to initial 120s to the following illuminations: LED (Dark red light) ND25 (green line, n= 4 cells), mCherry 100% (Green light) ND100 (blue line, n= 3 cells), or mCherry 25% (Green light) ND100 (orange line, n= 2 cells). Traction force listed as a function of initial force prior to illumination. Time t=0 represents the force at the instant the illumination period started. For both mCherry light sources, the cells keep relaxing for 120s until the illumination is over; after that they maintain the reduced traction for the next hour. LED light, on the other hand, did not elicit any immediate response which indicates that these cells are not affected by dark red light. Error bars represents standard error of the mean (SEM).

All of the studies above were limited to cells cultured on 2D substrates. To investigate the effect of light on cells in 3D extracellular matrix, we cultured NIH/3T3 fibroblasts in collagen matrix (rat-tail collagen I, Corning). After 1 hr of polymerization of the cell-ECM mixture, the cells were exposed to i) mCherry 100% ND25 (*λ* = 545-580 nm, I=7504 W/m^2^) ii) LED ND25 (*λ* = 635-650 nm, I=57 W/m^2^) illuminations for 120 sec. Following the exposure, we observed the cells for the next 10 hrs using only phase-contrast imaging. In the beginning, healthy control cells typically start to probe the microenvironment using filopodia and begin to elongate and migrate (Suppl. Vid. 1, n=10 cells). In stark contrast, cells exposed to mCherry 100% ND25 illumination underwent slowing down of activity followed by severe blebbing (Suppl. Vid. 2, n=10 cells) and apparent death. However, cells exposed to LED ND25 illumination appeared to be similar to control cells, i.e., they were elongated, contractile and migratory (Suppl. Vid. 3, n=10 cells). Hence, these results suggest that the threshold intensity of 57 W/m^2^ is possibly valid for NIH/3T3 fibroblasts within 3D collagen matrices as well.

## 4. Summary

Fluorescent excitation light has long been used to observe living cells, but little is known about the effect of light on cell functions. Currently, there exists no quantitative means to assess cell photo-response in real time. Furthermore, an exposure limit for mitigating photo-induced cell changes has not yet been established. Here, we search for a light intensity that has minimal effect on cells and yet that is sufficient for time-lapse fluorescent imaging. We exploit photo-sensitivity of fibroblasts to establish the threshold. Fibroblasts relax their contractility when exposed to light, and their photo-relaxation depends on light intensity, wavelength and exposure times. - We find that monkey (kidney fibroblast, CV-1), human (CCD-112 CoN, colon normal fibroblast) and mouse cells (NIH/3T3 fibroblasts) are almost insensitive to red light of wavelength, *λ*=635-650 nm and intensity, *I*~57 W/m^2^, even when they are subject to 60 s of continuous illumination. This insensitivity is independent of their extra cellular matrix. NIH/3T3 fibroblasts in 3D culture also show insensitivity to the same threshold. Furthermore, this light is sufficient for fluorescent imaging using standard fluorescent cameras. We thus suggest the use of *λ*> 650 nm with intensity 57 W/m^2^ as the potential light source for time-lapse imaging of living cells in 2D and 3D culture.

## Supporting information

Suppl. Vid. 1

Suppl. Vid. 2

Suppl. Vid. 3

Suppl. Vid. legends

## Acknowledgement

Research reported in this publication was supported by the National Institute of Biomedical Imaging and Bioengineering of the National Institutes of Health under Award Number T32EB019944 and NSF CMMI 17-42908. The content is solely the responsibility of the authors and does not necessarily represent the official views of the National Institutes of Health.

## Author Contributions

Conceived and designed the experiments: SK, BE, TS. Performed the experiments: BE, SK, UD, DB, LL, MS. Analyzed the data: BE, SK, UD, TS. Manuscript preparation: BE, SK, TS. All authors have read and approved the final manuscript.

## Disclosures

The authors declare no competing interests.

## Data Availability

All data generated or analyzed during this study are included in this published article (and its Supplementary Information files).

## Notes

https://drive.google.com/open?id=1PRs5Kma2t362DaYsf7uPXu8ExFAO9OMt

## References

1. Khodjakov, A. & Rieder, C. L. Imaging the division process in living tissue culture cells. Methods 38, 2–16 (2006).

2. Xiao, J. Single-Molecule Imaging in Live Cells. in Handbook of Single-Molecule Biophysics 43–93 (Springer US, 2009). doi:10.1007/978-0-387-76497-9_3

3. Furchgott, R. F. The Pharmacology of Vascular Smooth Muscle. Pharmacol. Rev. 7, (1955).

4. Ehrreich, S. J. & Furchgott, R. F. Relaxation of mammalian smooth muscles by visible and ultraviolet radiation. Nature 218, 682–4 (1968).

5. Knoll, S. G. Exposure to fluorescent excitation light induces dose-dependent, irreversible force relaxation in living fibroblast cells. (University of Illinois, Urbana-Champaign, 2016).

6. Stern, R. S. et al. Cutaneous Squamous-Cell Carcinoma in Patients Treated with PUVA. N. Engl. J. Med. 310, 1156–1161 (1984).

7. Kessel, D. H., Price, M., Reiners, J. J. & Jr. ATG7 deficiency suppresses apoptosis and cell death induced by lysosomal photodamage. Autophagy 8, 1333–41 (2012).

8. Lindl, T. & Steubing, R. Atlas of living cell cultures. (Wiley-Blackwell, 2013).

9. Lange, J. R. & Fabry, B. Cell and tissue mechanics in cell migration. Exp. Cell Res. 319, 2418–23 (2013).

10. Canović, E. P., Zollinger, A. J., Tam, S. N., Smith, M. L. & Stamenović, D. Tensional homeostasis in endothelial cells is a multicellular phenomenon. Am. J. Physiol. Physiol. 311, C528–C535 (2016).

11. Bellas, E. & Chen, C. S. Forms, forces, and stem cell fate. Curr. Opin. Cell Biol. 31, 92–7 (2014).

12. Taylor-Weiner, H., Ravi, N. & Engler, A. J. Traction forces mediated by integrin signaling are necessary for definitive endoderm specification. J. Cell Sci. 128, 1961–8 (2015).

13. Li, B. & Wang, J. H.-C. Fibroblasts and myofibroblasts in wound healing: force generation and measurement. J. Tissue Viability 20, 108–20 (2011).

14. Wang, J. H.-C. & Li, B. Application of Cell Traction Force Microscopy for Cell Biology Research. in 301–313 (Humana Press, 2009). doi:10.1007/978-1-60761-376-3_17

15. Emon, B., Bauer, J., Jain, Y., Jung, B. & Saif, T. Biophysics of Tumor Microenvironment and Cancer Metastasis - A Mini Review. Comput. Struct. Biotechnol. J. (2018). doi:10.1016/J.CSBJ.2018.07.003

16. Bauer, J. et al. Abstract 177: Increased stiffness of the tumor microenvironment in colon cancer leads to an increase in activin and metastatic potential. in Tumor Biology 78, 177–177 (American Association for Cancer Research, 2018).

17. Knoll, S. G., Ahmed, W. W. & Saif, T. A. Contractile dynamics change before morphological cues during fluorescence illumination. Sci. Rep. 5, 18513 (2016).

18. Brill-Karniely, Y. et al. Dynamics of cell area and force during spreading. Biophys. J. 107, L37–L40 (2014).

19. Nisenholz, N. et al. Active mechanics and dynamics of cell spreading on elastic substrates. Soft Matter 10, 7234 (2014).

20. Knoll, S. G. & Saif, M. T. A. Light induced, localized, and abrupt force relaxations in fibroblast cells on soft substrates. Extrem. Mech. Lett. 8, 257–265 (2016).

21. Knoll, S. G., Ali, M. Y. & Saif, M. T. A. A novel method for localizing reporter fluorescent beads near the cell culture surface for traction force microscopy. J. Vis. Exp. 51873 (2014). doi:10.3791/51873

22. Tse, J. R. & Engler, A. J. Preparation of Hydrogel Substrates with Tunable Mechanical Properties. Curr. Protoc. Cell Biol. 47, 10.16.1-10.16.16 (2010).

23. Iskratsch, T., Wolfenson, H. & Sheetz, M. P. Appreciating force and shape — the rise of mechanotransduction in cell biology. Nat. Rev. Mol. Cell Biol. 15, 825–833 (2014).

24. Roca-Cusachs, P., Iskratsch, T. & Sheetz, M. P. Finding the weakest link – exploring integrin-mediated mechanical molecular pathways. J. Cell Sci. 125, 3025–3038 (2012).

25. Reinhart-King, C. A., Dembo, M. & Hammer, D. A. The Dynamics and Mechanics of Endothelial Cell Spreading. Biophys. J. 89, 676–689 (2005).

26. Saez, A., Buguin, A., Silberzan, P. & Ladoux, B. Is the Mechanical Activity of Epithelial Cells Controlled by Deformations or Forces? Biophys. J. 89, L52–L54 (2005).

27. Wäldchen, S., Lehmann, J., Klein, T., van de Linde, S. & Sauer, M. Light-induced cell damage in live-cell super-resolution microscopy. Sci. Rep. 5, 15348 (2015).

