## Supplementary material for "Threshold illumination for non-invasive imaging of cells and tissues": Suppl. Vid. legends

**Suppl. Video 1: Control 3T3 fibroblast cells, not exposed to any monochromatic lights, exhibit normal activities in 3D collagen matrix.** Observation started after 1 hr of collagen polymerization. The cell was imaged every 5 mins with only phase-contrast illumination for 10 hrs and it looks active, contractile and motile throughout. The whole video represents 10 hrs at 7 fps.

**Suppl. Video 2: 3T3 fibroblasts exposed to green light (above threshold) for 120s initially, embraces apparent death within hours in 3D collagen matrix.** After 1 hr of collagen polymerization, the cells were exposed to mCherry 100% ND25 (λ=545-580 nm, I=7504 W/m^2^) illumination for 120 s and observation followed. Images were taken every 5 mins with only phase-contrast illumination for 10 hrs. Initially the cell appears to be healthy and active; but within 4-6 hours, starts to show signs of distress, excessive blebbing and possible death. The whole video represents 10 hrs at 7 fps.

**Suppl. Video 3: 3T3 fibroblasts exposed to red light (below threshold) for 120s initially, shows normal behavior in 3D collagen matrix.** After 1 hr of collagen polymerization, the cells were exposed to LED ND25 (λ=635-650 nm, I=57 W/m^2^) illumination for 120 s and then observation followed. The cell was imaged every 5 mins with only phase-contrast light for 10 hrs and it seems to be healthy, contractile and migratory as the control cell. The whole video represents 10 hrs at 7 fps.
