## Supplementary material for "Threshold illumination for non-invasive imaging of cells and tissues": Suppl. information

### Supplementary Information:

Fluorescent Light source:

X-Cite® Series 120Q (Excelitas Technologies, Waltham, MA) coupled with an mCherry filter (Semrock Brightline mCherry-M-OMF, λ =540-585/600-682 nm ex/em, Rochester, NY) was used for green light illumination. X-Cite controller can regulate the power of the light (100%, 50%, 25% and 12%). Further reduction in intensity can be achieved by using ND filters. Suppl. Fig. 1a shows the spectrum of light for 25%mCherry ND25.

LED light source:

For red illumination, we used an LED light source (Model M660L3-C1, S/N M00297422, Thorlabs, Inc.). This LED was coupled with a collimation adapter (COP1-A, AR Coating: 350-700 nm) and was controlled by a LEDD1B T-Cube LED Driver. The driver can control both the current and power to modulate the intensity of light. We also added a Semrock BrightLine long-pass laser filter set (LF635/LP-B-000) for blocking out any light with lower wavelength. Suppl. Fig. 1b shows the spectrum of light for LED ND25.

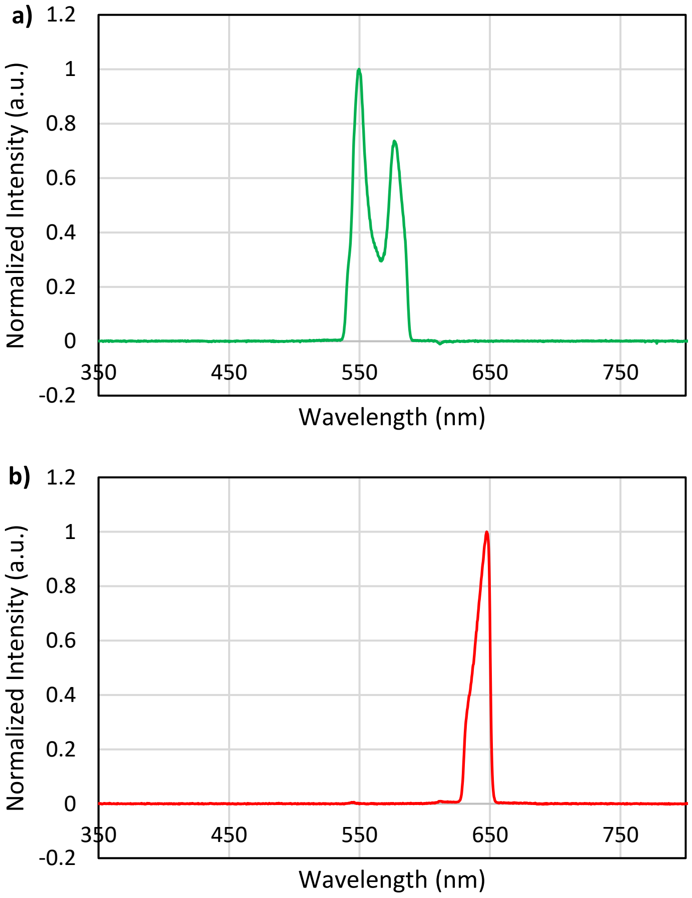


**Suppl. Fig. 1**: Light spectrum for excitation lights a) mCherry b) LED

Experimental setup:

Suppl. Fig. 2 presents a simplified schematic diagram of the experimental setup. The microscope we used is an Olympus IX81 with an environment control chamber. The cells were maintained at 37 degree Celsius and 5% $CO_2$ while the experiments were running. The brightfield light illumination is from the top of the sample and is collected through the objective in the image acquisition system. On the other hand, the LED and fluorescent lights come through the objective and illuminate the sample from the bottom. The emitted light goes back into the objective and then collected in the acquisition system. The schematic shows the location of the filter sets, shutter and ND filters along the light path. A manual switch controls whether light from the LED or X-Cite is going to be used for illumination. The bottom shutter is program controlled and can be used for sub-second exposure experiments. It can be noted that we could switch between LED and X-Cite light sources manually whenever necessary. This enabled us to illuminate with one light, and then observe the beads with another light. The brightfield light illumination is from the top, and thus it does not interfere with the LED or X-Cite light paths.

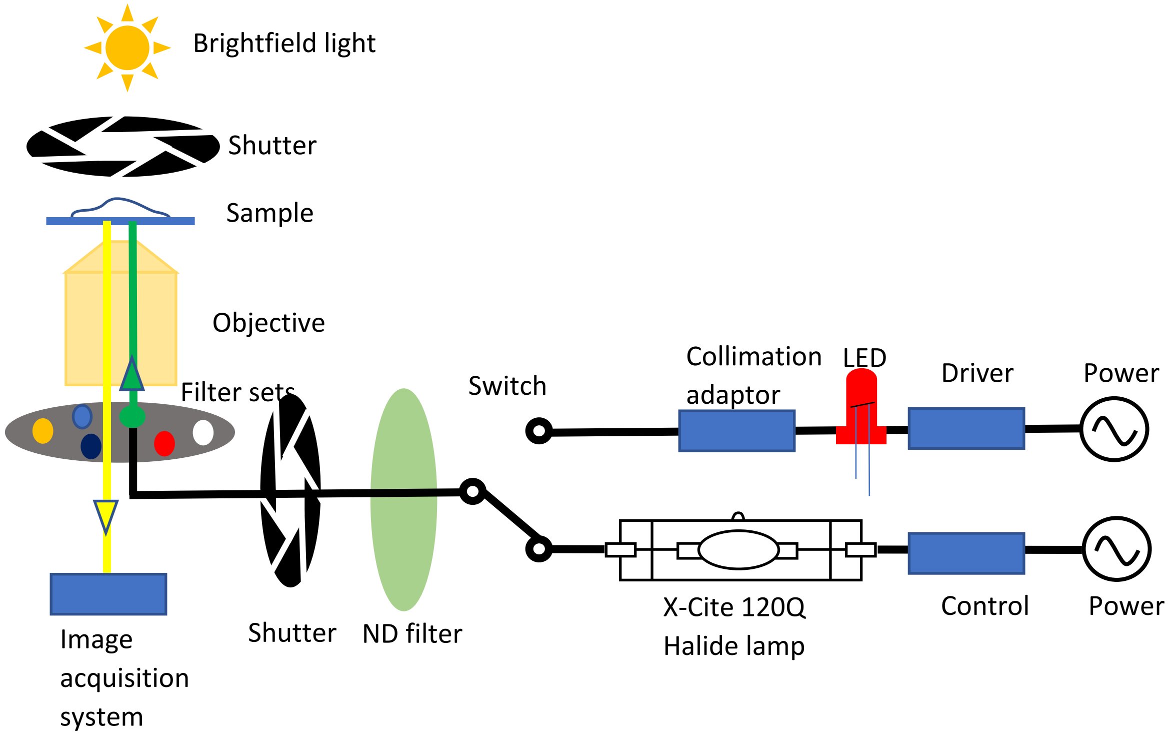


**Suppl. Fig. 2**: Experimental setup for illumination

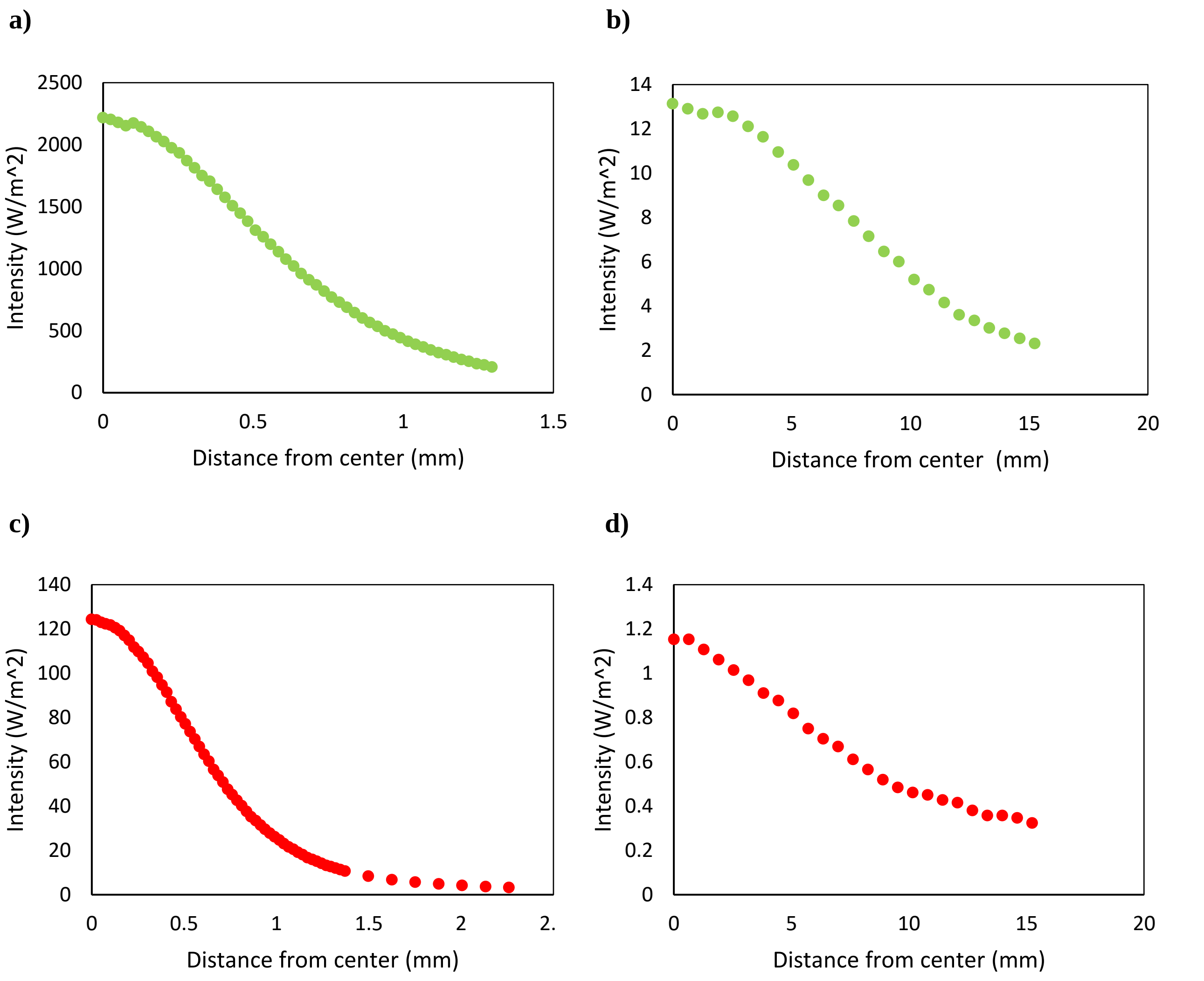


**Suppl. Fig. 3. Intensity profile of light beam through 40X water immersion objective for different light sources at 15.5 mm from the lens and focal plane (FP).** Light intensity was measured using PM100 power meter (Thorlabs). Light sources are a) mCherry25%ND25 at FP b) mCherry25%ND25 at 15.5 mm c) LED ND25 at FP d) LED ND25 at 15.5 mm.

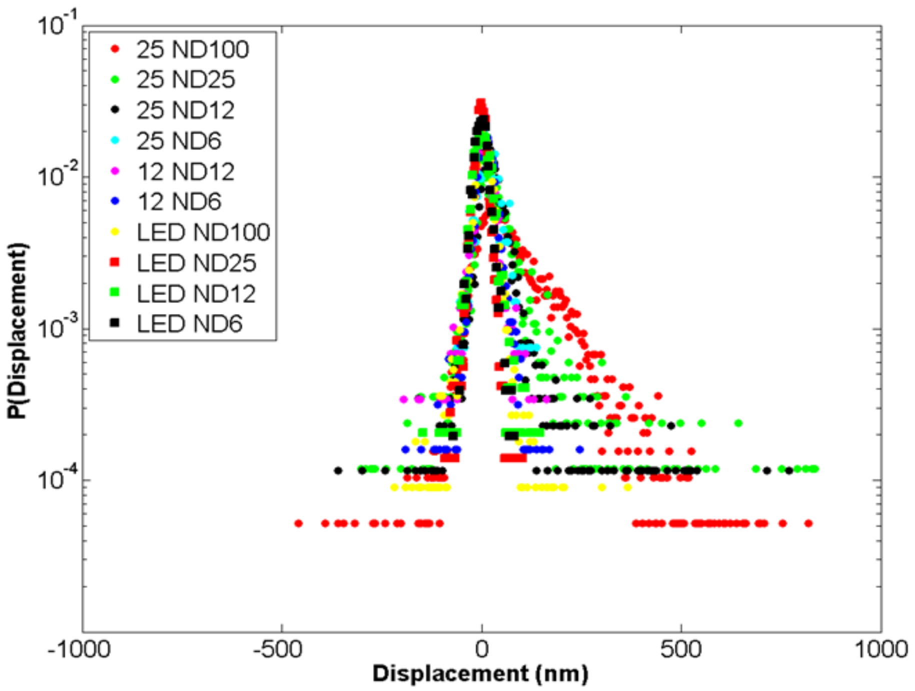


**Suppl. Fig. 4. Probability of positive displacements decreases with decreasing light intensity.** Displacement distributions for cells exposed to various light sources. All cells were illuminated continuously for 60 s. Each distribution represents n=5 distinct cells.

**a)**

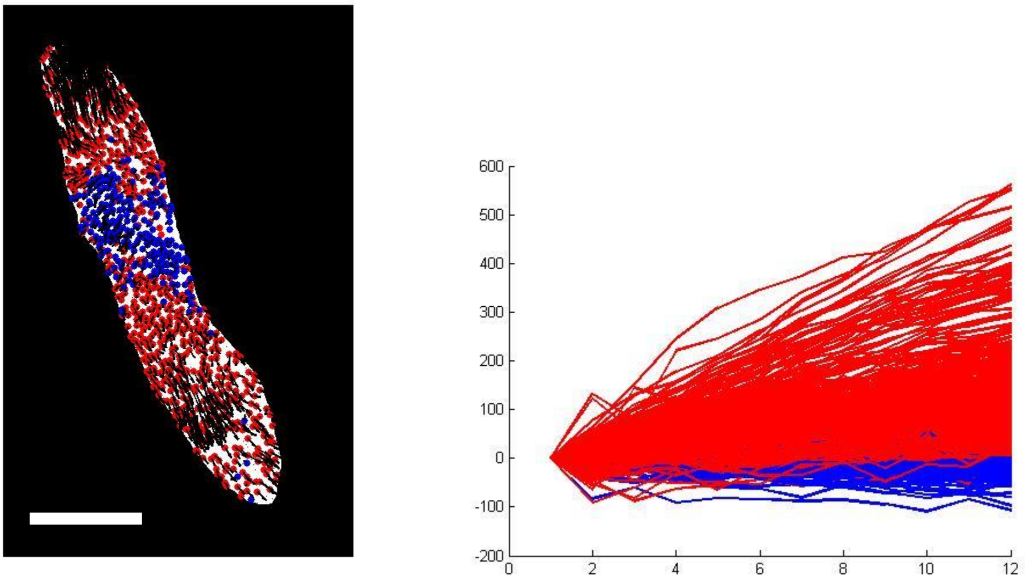


**b)**

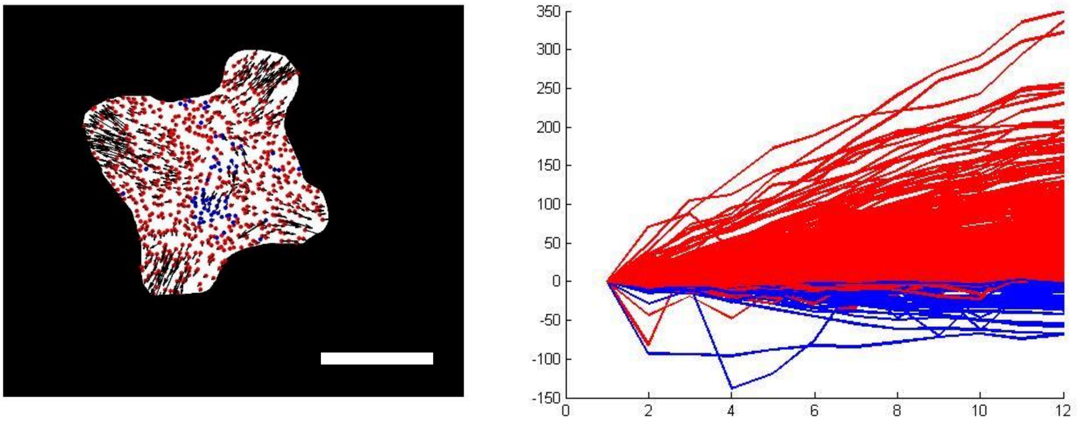


**Suppl. Fig. 5. Displacements of beads under cells cultured on 5 Kpa PA gel substrate functionalized with laminin.** The cells are subjected to a)100% mCherry (green) ND25, 120s and b) 100% mCherry (green) ND100, 120s. Red and blue trajectories indicate relaxation (away from cell center) and contraction (towards cell center) respectively. Most of the beads continue to relax for 120 s during the light exposure. These results are similar to those with fibronectin ECM, implying that photo relaxation is independent of ECM. Scale bars: 20 um.

Dosage calculation:

For experiments conducted for the study, calculated initial dosage and cumulative dosage due to observation is presented in Table S1. For most of the experiments, initial target dosage is more than 96% of the total dosage which indicates that the effect of observation should be minimal.

Table S1:

| **Figure number** | **Target Illumination** | **Exposure duration** | **Exposure dosage (J/m$^2$)** | **Observation period** | **Observation illumination** | **Observation frequency** | **Imaging exposure, Dosage (J/m$^2$)** | **Total dosage (J/m$^2$)** | **Target dosage percentage** |
|---|---|---|---|---|---|---|---|---|---|
| 1A, 2 | 25%mCherry ND100 | 60 s | 441063 | 60 s | 25%mCherry ND100 | - | - | 441063 | 100% |
| 1B, 2 | 25%mCherry ND25 | 60 s | 135130 | 60 s | 25%mCherry ND25 | - | - | 135130 | 100% |
| 1C, 2 | 25%mCherry ND12 | 60 s | 59781 | 60 s | 25%mCherry ND12 | - | - | 59781 | 100% |
| 1D, 2 | 25%mCherry ND6 | 60 s | 26702 | 60 s | 25%mCherry ND6 | - | - | 26702 | 100% |
| 1E, 2 | 12%mCherry ND12 | 60 s | 34269 | 60 s | 12%mCherry ND12 | - | - | 34269 | 100% |
| 1F, 2 | 12%mCherry ND6 | 60 s | 15459 | 60 s | 12%mCherry ND6 | - | - | 15459 | 100% |
| 1G, 2 | LED ND100 | 60 s | 14087 | 60 s | LED ND100 | - | - | 14087 | 100% |
| 1H, 2 | LED ND25 | 60 s | 3430 | 60 s | LED ND25 | - | - | 3430 | 100% |
| 1I, 2 | LED ND12 | 60 s | 2044 | 60 s | LED ND12 | - | - | 2044 | 100% |
| 1J, 2 | LED ND6 | 60 s | 876 | 60 s | LED ND6 | - | - | 876 | 100% |
| 3 | LED ND100 | 2 s | 470 | 60 min | LED ND25 | every 5 mins | 0.2s, 11.44 | 607 | 77% |
| 3 | LED ND100 | 15 s | 3522 | 60 min | LED ND25 | every 5 mins | 0.2s, 11.44 | 3659 | 96% |
| 3 | 25%mCherry ND25 | 2 s | 4504 | 60 min | LED ND25 | every 5 mins | 0.2s, 11.44 | 4641 | 97% |
| 3 | 25%mCherry ND25 | 15 s | 33783 | 60 min | LED ND25 | every 5 mins | 0.2s, 11.44 | 33920 | 100% |
| 4 | LED ND100 | 2 s | 470 | 60 min | LED ND25 | 10, 30 & 60 min | 0.2s, 11.44 | 504 | 93% |
| 4 | LED ND100 | 15 s | 3522 | 60 min | LED ND25 | 10, 30 & 60 min | 0.2s, 11.44 | 3556 | 99% |
| 4 | 25%mCherry ND100 | 2 s | 14702 | 60 min | LED ND25 | 10, 30 & 60 min | 0.2s, 11.44 | 14736 | 100% |
| 5 | LED ND100 | 2 s | 470 | 60 min | LED ND25 | 10, 30 & 60 min | 0.2s, 11.44 | 504 | 93% |
| 5 | 25%mCherry ND100 | 15 s | 110267 | 60 min | LED ND25 | 10, 30 & 60 min | 0.2s, 11.44 | 110301 | 100% |
| 6 | 100%mCherry ND100 | 60 s | 1469622 | 60 s | 100%mCherry ND100 | - | - | 1469656 | 100% |
| 6 | 100%mCherry ND25 | 60 s | 450252 | 60 s | 100%mCherry ND25 | - | - | 450286 | 100% |
| 7 | 25%mCherry ND100 | 1 s | 7351 | 60 min | LED ND25 | every 5 mins | 0.2s, 11.44 | 7488 | 98% |
| 7 | 25%mCherry ND100 | 500 ms | 3676 | 60 min | LED ND25 | every 5 mins | 0.2s, 11.44 | 3813 | 96% |
| 8 | LED ND25 | 120 s | 6864 | 60 min | LED ND25 | 20, 40 & 60 min | 0.2s, 11.44 | 6898 | 100% |
| 8 | 25%mCherry ND100 | 120 s | 882132 | 60 min | LED ND25 | 20, 40 & 60 min | 0.2s, 11.44 | 882166 | 100% |
| 8 | 100%mCherry ND100 | 120 s | 2939244 | 60 min | LED ND25 | 20, 40 & 60 min | 0.2s, 11.44 | 2939278 | 100% |
